# Endosomal GLUT3 is essential for alternative macrophage signaling, polarization, and function

**DOI:** 10.1101/2022.11.09.515804

**Authors:** Dong-Min Yu, Jiawei Zhao, Eunice E. Lee, Elysha K. Rose, Ruchika Mahapatra, Jun-Yong Choe, E Dale Abel, Richard C. Wang

## Abstract

Macrophages play critical roles in both inflammation and tissue homeostasis. Classically activated (M1) macrophages promote antimicrobial and tumoricidal activity, while alternatively activated (M2) macrophages promote phagocytosis and tissue homeostasis. The facilitative GLUT1 and GLUT3 hexose transporters are expressed abundantly in different hematopoietic lineages, but their specific functions in macrophages is poorly understood. We discovered that GLUT3 expression was increased after M2-activation stimuli in macrophages. Notably, GLUT3 KO BMDM (bone marrow-derived macrophages) showed marked defects in M2, but not M1, polarization. Consistent with defects in M2 polarization, GLUT3 KO macrophages showed impaired wound healing and decreased inflammation in calcipotriol-induced, atopic dermatitislike inflammation. GLUT3 promoted IL-4/STAT6 signaling, the main signaling pathway for M2 polarization, in a glucose-transport independent manner. Unlike plasma membrane-localized GLUT1, GLUT3 and components of the IL-4 signaling pathway, localized primarily to endosomes. GLUT3, but not GLUT1, interacted with Ras through its intracytoplasmic loop, and Rac1-PAK-cofilin signaling and the endocytosis of IL4R subunits were impaired in the absence of GLUT3. Thus, GLUT3 is essential for alternative macrophage polarization and function and plays an unexpected role in the regulation of endosomal signaling.

## INTRODUCTION

Macrophages are immune cells that play critical roles in both inflammation and tissue homeostasis. While macrophages appear to exhibit dynamic and complex functional phenotypes in vivo, the dichotomous model of macrophage activation remains a critical paradigm to understand macrophage functions (Mantovani et al., 2005; Munoz-Rojas et al., 2021). Classically activated (M1) macrophages typically promote antimicrobial and tumoricidal activity, while alternatively activated (M2) macrophages promote phagocytosis and tissue homeostasis (Ley, 2017; Sica and Mantovani, 2012). M1 polarization can be induced by Interferon gamma (IFN-γ) and Toll-Like Receptor (TLR) agonists such as lipopolysaccharides (LPS), while M2 polarization is induced by IL-4 or IL-13 (Murray, 2017; Orecchioni et al., 2019). During the M1 polarization process, Nuclear Factor kappa-light-chain enhancer of activated B cell (NF-κB) and Signal Transducer and Activator of Transcription 1 (STAT1) are activated, while M2 polarization is mainly regulated by the activation of STAT6, which then result in the expression and function of M1 and M2-specific markers. For the STAT6 signaling pathway, the binding of ligands to receptors leads to the activation of Janus kinases (JAK). Activated JAK phosphorylates receptor tyrosine residues, and phospho-tyrosine sites of receptor serve as docking sites for STAT6(Hu et al., 2021).

Glucose transporters are responsible for the first step of glucose utilization in cells, and 14 facilitative glucose transporters (GLUTs) are expressed in humans (Navale and Paranjape, 2016). Among them, GLUT1 and GLUT3 were found to be expressed in human lymphocytes and macrophages (Fu et al., 2004). Both GLUT1 and GLUT3 are class I glucose transporters, and despite high similarity in amino acid sequence and structure, the two transporters show differences in their pattern of tissue expression and expression levels in different cell types (Deng et al., 2015; Deng et al., 2014). GLUT1, the most widely expressed facilitative glucose transporter, is highly expressed in erythrocytes, blood-brain barrier endothelial cells, and keratinocytes (Cura and Carruthers, 2012; Zhang et al., 2018). GLUT3 shows a more tissuespecific expression pattern than GLUT1 and is highly expressed in neurons and hematopoietic lineage cells (Fidler et al., 2017; Simpson et al., 2008). Functional studies examining the isoform specific functions of GLUTs in macrophages, and leukocytes in general, remain limited.

Endocytosis has traditionally been known as a mechanism to prevent excessive ligand-induced activation of downstream effectors by removing activated receptors on the cell surface (Sorkin and von Zastrow, 2009). However, endosomes can also act as a signaling platform for numerous receptor tyrosine kinases (RTKs), including epidermal growth factor receptor (EGFR), by ensuring sufficient duration and intensity of signaling (Grimes et al., 1996; Vieira et al., 1996) In particular, for IL-4 receptor signaling, endosomes have been found to be essential for efficient ligand-induced receptor dimerization and signal transduction. IL-4 receptor subunits endocytosis is distinct from the endocytosis of many RTKs and transforming growth factor beta (TGF-β) and has been found to be Rac1-, Pak- and actin-mediated (Kurgonaite et al., 2015).

In this study, we evaluated the subcellular localization and function of the most highly expressed glucose transporters in macrophages, GLUT1 and GLUT3. We confirmed that GLUT1 expression was increased in M1 macrophages and discovered that GLUT3 expression was increased in M2 macrophages. Notably, GLUT3 KO BMDM (bone marrow-derived macrophages) showed a defect in M2 polarization *in vitro* and *in vivo*. IL-4-STAT6 signaling, the main signaling for M2 polarization, was impaired by GLUT3 deficiency. Unlike GLUT1, which localized to the plasma membrane, GLUT3, along with components of the IL-4 signaling pathway, localized to endosomes. Finally, we found that GLUT3 interacts with Ras and regulates the Rac1-PAK-actin pathway, which regulates IL-4 receptor endocytosis. Thus, our studies reveal that endosomal GLUT3 is essential for M2 polarization of macrophages by regulating IL-4-STAT6 signaling.

## RESULTS

### GLUT3 is induced by M2 stimuli and required for M2 polarization of macrophages

To investigate the specific functions of GLUT isoforms in macrophages, we began by determining their expression levels after polarization stimuli. Mouse bone marrow-derived macrophages (BMDMs) were treated with lipopolysaccharide (LPS) and interferon gamma (IFN-g) to induce M1 polarization, and IL-4 to induce M2 polarization, and expression of facilitative (GLUT) and sodium dependent (SGLT) glucose transporters isoforms were assessed by quantitative real-time RT-PCR (qRT-PCR). Consistent with previous studies, GLUT1 mRNA expression was elevated in M1 macrophages (Cho et al., 2022; Freemerman et al., 2019). Notably, GLUT3 mRNA expression was significantly elevated in M2 macrophages (Fig. 1a). This finding was reproduced in additional macrophages cell lines, including human THP-1 cells and murine RAW 264.7 cells. GLUT1 and GLUT3 mRNA expression were similarly increased by M1 and M2 stimuli, respectively (Fig. 1b-c). Given the striking regulation of GLUT1 and GLUT3 by polarization, we focused on these two transporters and their impact on macrophage polarization markers. We generated WT (*Slc2a1*^flox/flox^; LysM-Cre^-^ or *Slc2a3*^flox/flox^; LysM-Cre^-^) and myeloid cell-specific GLUT1 KO (*Slc2a1*^flox/flox^; LysM-Cre^+^) and GLUT3 KO (*Slc2a3*^flox/flox^; LysM-Cre^+^) BMDMs to address these questions. Consistent with previous reports, we found that GLUT1 KO BMDMs showed defects in M1 polarization as shown by the strong reduction in common M1 markers such as Nos2 (nitric oxide synthase 2), Tnfa (tumor necrosis factor a) and Il1b (interleukin-1 b). No significant difference in M2 markers after M2 polarization were noted between WT and GLUT1 KO BMDMs (Fig. 1d). Strikingly, GLUT3 KO BMDMs showed an increased expression of M1 markers compared to WT after M1 polarization, whereas the expression of common M2 markers Arg1 (arginase), Retnla (resistin like alpha), and Chil3l3 (chitinase-like 3) was significantly reduced after M2 polarization (Fig. 1e). We next investigated the effect of GLUT 1 or GLUT3 deficiency on glucose uptake assays using 2-deoxyglucose uptake assays in macrophages. Glucose uptake was decreased in GLUT1 KO BMDMs compared to WT BMDMs by M1 stimuli, but there was no difference in glucose uptake after M2 stimuli. Consistent with the increased expression of M1 polarization markers, GLUT3 KO BMDMs showed increased glucose uptake after M1 stimuli compared to WT BMDMs. However, there was no difference in 2-DG uptake from the media after M2 polarization (Fig. 1f).

**Figure 1.**
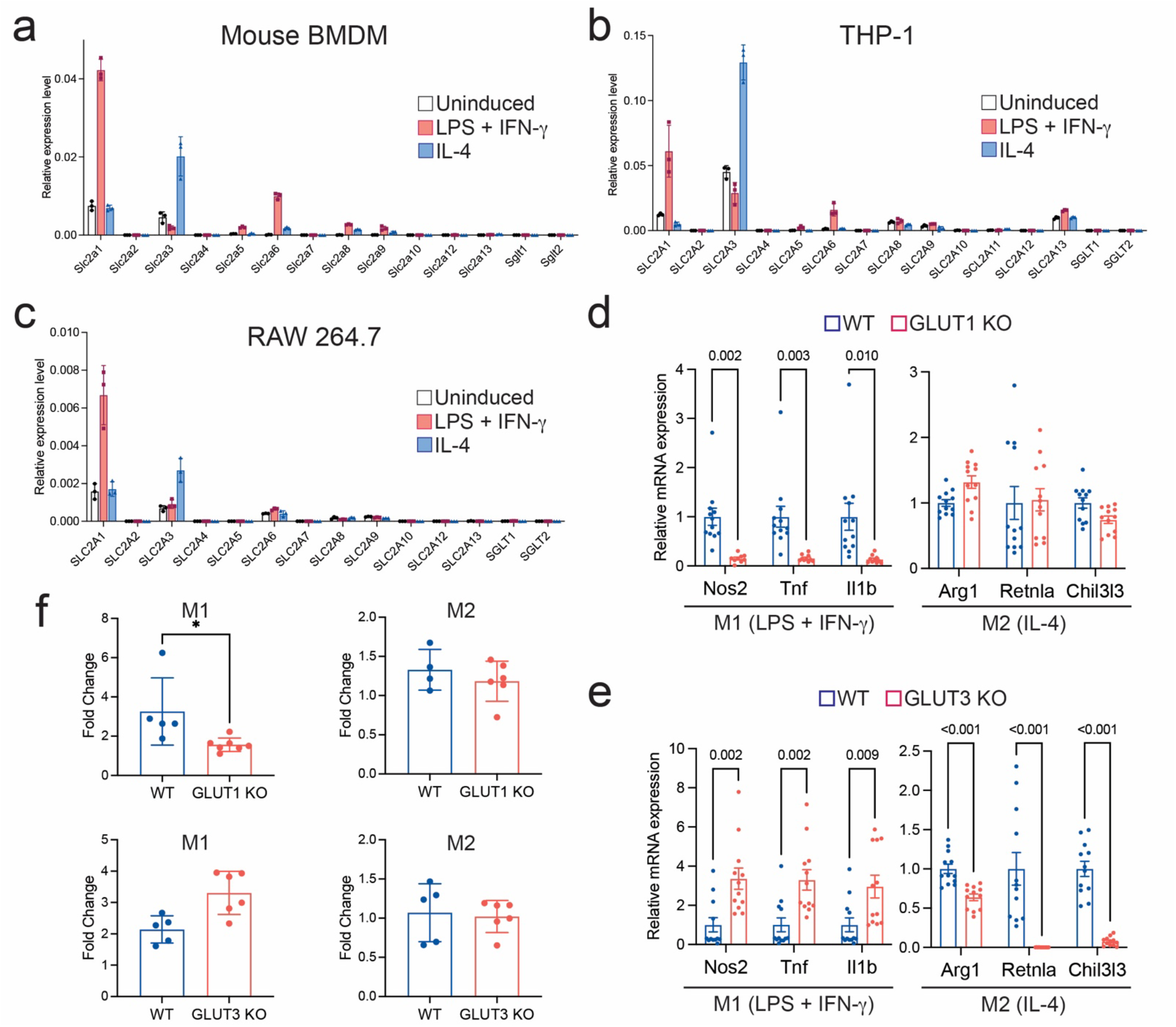
Expression of GLUT3 is increased by M2 stimulus, and GLUT3 deficiency impairs M2 polarization of macrophages. **(a-c)** mRNA expression levels of GLUT and SGLT transporter isoforms in BMDMs (a), THP-1 cells (b), and Raw 264.7 cells (c) in unstimulated macrophages (white) and after treatment with classic M1 (red) or alternative M2 (blue) polarization stimuli for 24 hours. Expression normalized to β-actin (ACTB) expression **(d)** mRNA expression levels of M1 (Nos2, Tnfa and Il1b) and M2 (Arg1, Retnla and Chil3l3) markers in WT (n=12) and GLUT1 KO (n=12) BMDMs after the indicated polarization stimuli. **(e)** mRNA expression levels of M1 and M2 markers in WT (n=12) and GLUT3 KO (n=12) BMDMs after the indicated polarization stimuli. **(f)** 2-Deoxy-D-glucose uptake in WT, GLUT1 KO, and GLUT3 KO BMDMs after the indicated polarization stimuli. Data shown as mean ± SEM. P values were calculated by two-tailed t-test. *P < 0.05, **P < 0.01, ***P < 0.001.

### GLUT3 KO macrophages rescue a mouse model of calcipotriol induced atopic dermatitis

Given the striking impact of GLUT3 deficiency on M2 polarization in vitro, we next investigated its impact on macrophage function in vivo. While M2 macrophages are best known for their functions in tissue homeostasis, they have also been shown to play important roles in promoting allergic (Type 2) inflammation (Kasraie and Werfel, 2013; Suzuki et al., 2017). Thus, we studied the impact of GLUT3 deficiency in a mouse model of atopic dermatitis. Dermatitis was induced in WT, GLUT1, and GLUT3 KO mice through the topical administration of calcipotriol (MC903) and the development of the inflammation was assessed (Li et al., 2006; Oetjen et al., 2017) (Fig. 2a). The back and treated ears of WT mice and GLUT1 KO mice edema, erythema, and scaling, consistent with previous reports of the model. GLUT3 KO mice showed notably less inflammation with decreased edema, erythema, and scale compared to WT and GLUT1 KO mice (Fig. 2b). Consistent with the macroscopic observations, histological analyses revealed decreased epidermal hyperplasia and hyperkeratosis in GLUT3 KO mice compared to GLUT1 KO mice or WT mice (Fig. 2c-d). To extend these findings, calcipotriol-treated tissues were harvested (Day 13) and qRT-PCR was used to detect markers of macrophage polarization. F4/80, a pan-macrophage marker was consistent throughout the WT, GLUT3 and GLUT1 KO mice. In addition, there were no significant changes in M1 markers such as Nos2 and Tnfa between WT, GLUT3 and GLUT1 KO mice (Fig. 2e). However, there were significant reduction in M2 marker levels in GLUT3 KO mice compared to WT and GLUT1 KO mice (Fig. 2f). In addition, we found that levels of several Th2 cytokines, including IL-4, IL-13, and IL-31, which are known to increase in response to calcipotriol treatment, (Li *et al.*, 2006; Li et al., 2005) were significantly reduced after GLUT3 KO compared to WT mice (Fig. 2g).

**Figure 2.**
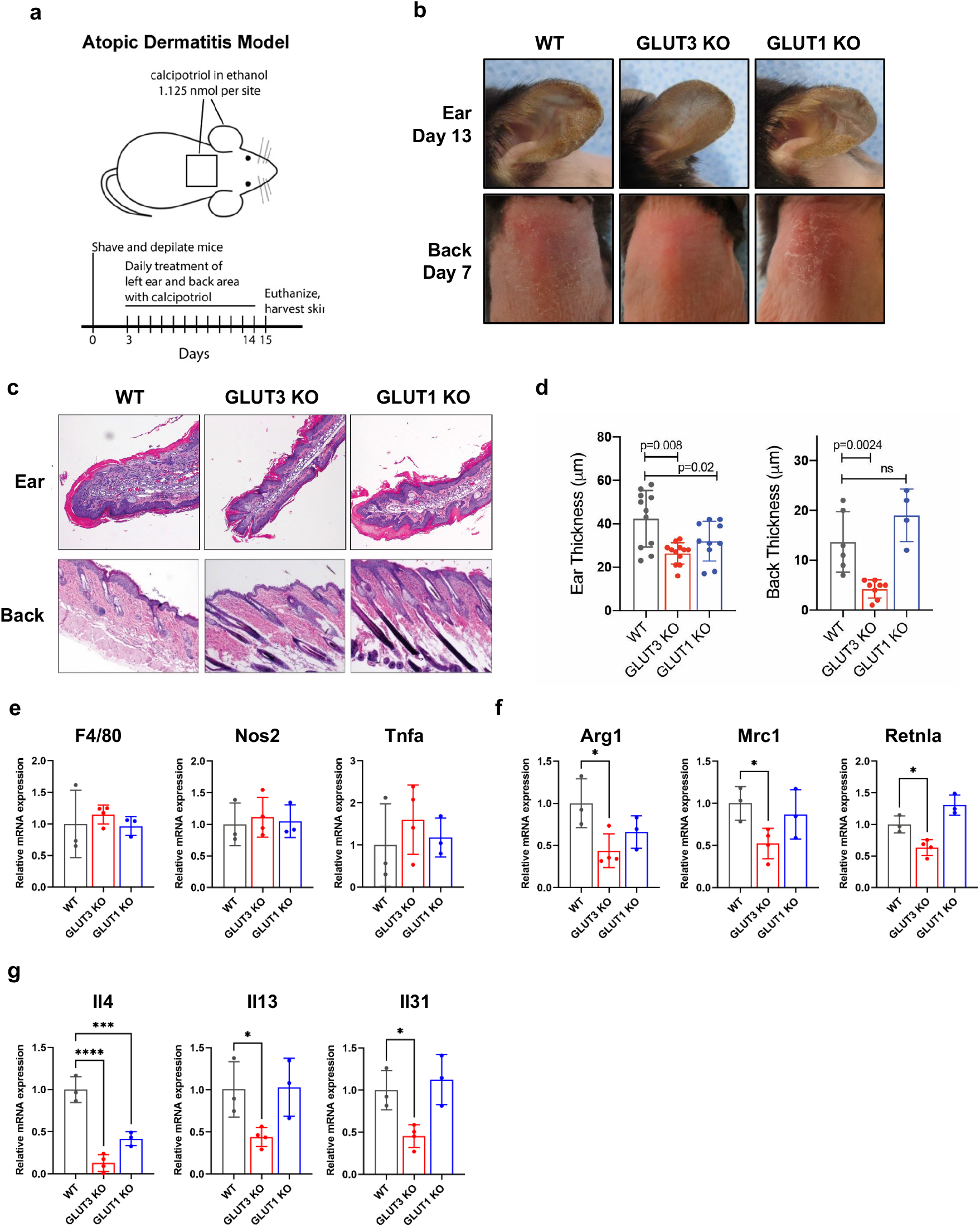
GLUT3 KO macrophages rescue a mouse model of calcipotriol induced atopic dermatitis like rash. **(a)** Scheme for calcipotriol (MC903) induced dermatitis. Calcipotriol (1.125 nmol) in ethanol was applied to the left ear and shaved back of the indicated mice for 13 days. **(b)** Representative photos after calcipotriol administration in WT, GLUT3 KO and GLUT1 KO mice at day 8. **(c)** Hematoxylin and eosin stained sections of mouse skin treated with calcipotriol analyzed at day 13 in WT, GLUT3 KO, and GLUT1 KO mice. **(d)** Thickness of calcipotriol-treated ear and back in WT (n=11 for ear and n=6 for back), GLUT3 KO (n=12 for ear and n=8 for back), and GLUT1 KO (n=10 for ear and n=4 for back) mice. **(e-g)** mRNA expression levels in calcipotriol-treated ear in WT (n=3), GLUT3 KO (n=4), and GLUT1 KO (n=3) mice. Pan-macrophage marker (F4/80), M1 markers (Nos2 and Tnfa) (e), M2 markers (Arg1, Mrc1 and Retnla) (f), and Th2 cytokines (IL-4, IL-13, and IL-31) (g) were observed. Data shown as mean ± SEM. P values were calculated by two-tailed t-test. *P < 0.05, **P < 0.01, ***P < 0.001.

### Wound healing is delayed in GLUT3 KO mice

M2 macrophages have also been shown to play a critical role in wound healing, particularly in the later phases of neovascularization and tissue remodeling (Rehak et al., 2022). Thus, ss a second *in vivo* mouse model, we performed a splinted, excisional wound healing model. The back skin of WT, GLUT1, and GLUT3 KO mice were excised by punch biopsy, splinted, and wound healing was measured every two days (Fig. 3a). In GLUT3 KO mice, wound healing was significantly delayed compared to WT and GLUT1 KO mice (Fig. 3b-c). We also examined macrophage markers from tissue obtained from the wound edge. Consistent with the atopic dermatitis model, there was no difference in the expression of total macrophage and the M1 macrophage markers, but the expression of the M2 markers was significantly reduced in GLUT3 KO mice (Fig. 3d-e). Immunofluorescence (IF) of the wound tissues was used to corroborate the qRT-PCR studies. Specifically, wound tissues were co-stained with the pan-macrophage marker, F4/80, and the M2 marker, Arg1 (arginase). While there was no significant difference in the degree of F4/80 staining in WT, GLUT1 KO, and GLUT3 KO mice, the expression of Arg1 (arginase1) and the number of F4/80^+^ Arg1^+^ macrophages was significantly reduced in GLUT3 KO mice (Fig. 3f-g). Finally, additional markers relevant to tissue remodeling and angiogenesis, which have previously been shown be involved in the wound healing process, were also consistently reduced in GLUT3 KO mice (Koo et al., 2019; Okonkwo et al., 2020; Zomer and Trentin, 2018) (Fig. 3h-i). Thus, using two in vivo models of M2 polarization, we conclude that myeloid cell-specific GLUT3 is required for M2 polarization and function.

**Figure 3.**
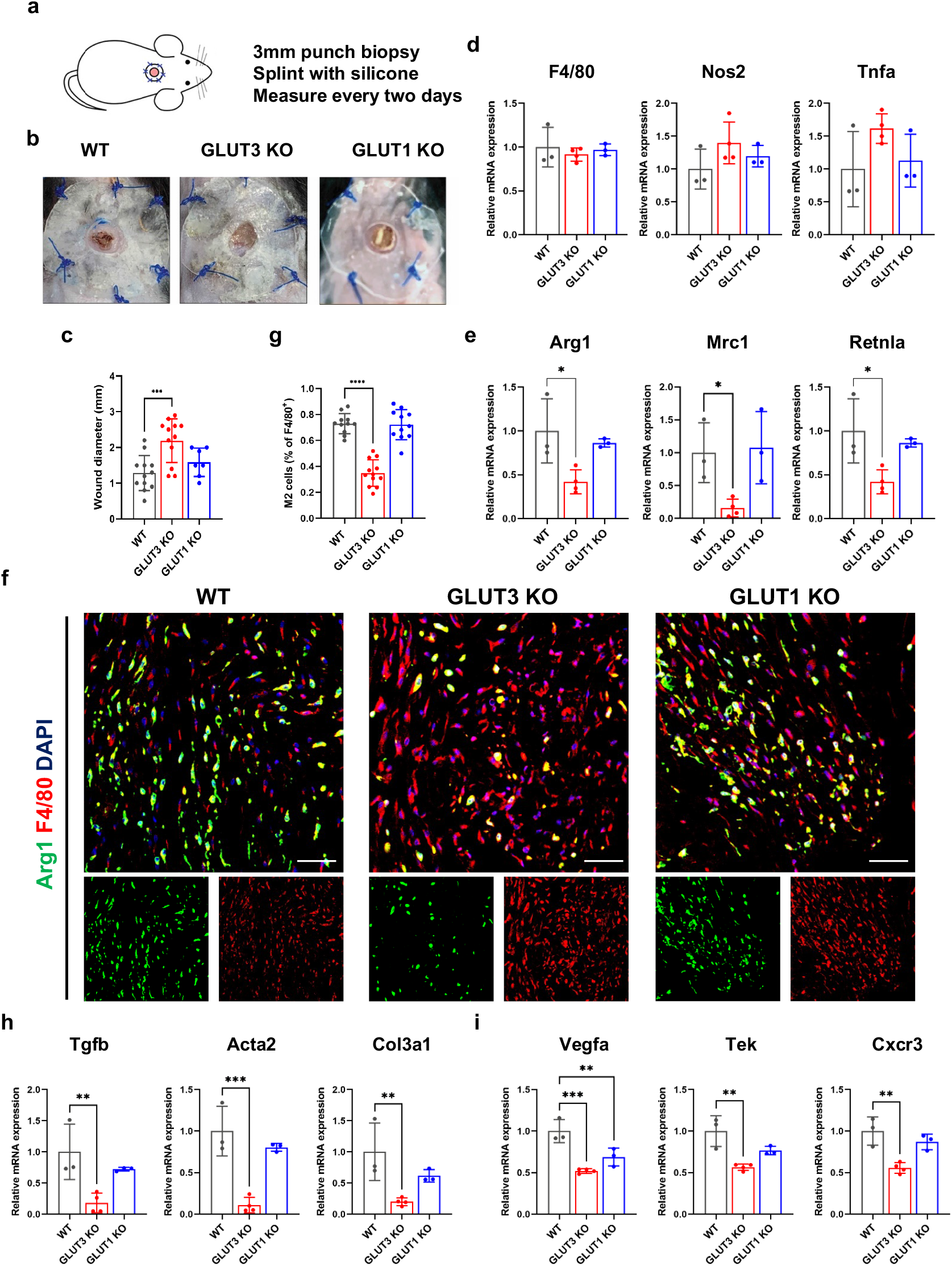
Delayed wound healing in GLUT3 KO mice. **(a)** Scheme for splinted wound healing model. Wounds on the shaved back of WT, GLUT1, and GLUT3 KO mice were generated by 3.0 mm punch biopsy, splinted, and wound healing was measured every two days. **(b)** Representative photos of wound site in WT, GLUT3 KO and GLUT1 KO mice at 6 days after injury. **(c)** Measurements of wound diameter on day 6 in WT (n=12), GLUT3 KO (n=12) and GLUT1 KO (n=7) mice. **(d-e)** mRNA expression levels of F4/80, Nos2 and Tnfa in WT (n=3), GLUT3 KO (n=3) and GLUT1 KO (n=3) mice. **(f)** Representative immunofluorescence stains of Arg1 (green), F4/80 (red) and DAPI (blue) in the wound site at 6 days after injury. Scale bar = 50 μm. (**g**) Statistical analysis of M2 macrophages at wound sites. The number of F4/80^+^Arg1^+^ cells was normalized to the number of F4/80^+^ cells. **(h-i)** mRNA expression levels of tissueremodeling related markers (Tgfb, Acta2 and Col3a1) (h) and angiogenesis markers (Vegfa, Tek and Cxcr3) (i). Data shown as mean ± SEM. P values were calculated by two-tailed t-test. *P < 0.05, **P < 0.01, ***P < 0.001.

### STAT6 signaling is impaired by GLUT3 deficiency

To elucidate the mechanism of the relationship between GLUT3 and M2 polarization in macrophages, we investigated macrophage polarization signaling in greater detail. There was no difference in phosphorylation of STAT1 by LPS and IFN-g in GLUT1 KO BMDMs, but phosphorylation of STAT6 by IL4 was notably reduced in GLUT3 KO BMDMs (Fig. 4a-b). The effects on STAT6 signaling were confirmed in human and mouse macrophage cells lines. GLUT3 knockdown also strongly reduced the expression of phospho-STAT6 in THP-1 and RAW264.7 cells (Fig. 4c-d). We next determined whether GLUT3 also affected phosphorylation of JAK1 upstream of STAT6 in IL4 signaling. Levels of phospho-JAK1 was also decreased by GLUT3 deficiency in BMDMs (Fig. 4e). These findings reveal that GLUT3 is required for M2 polarization by promoting IL4 signaling.

**Figure 4.**
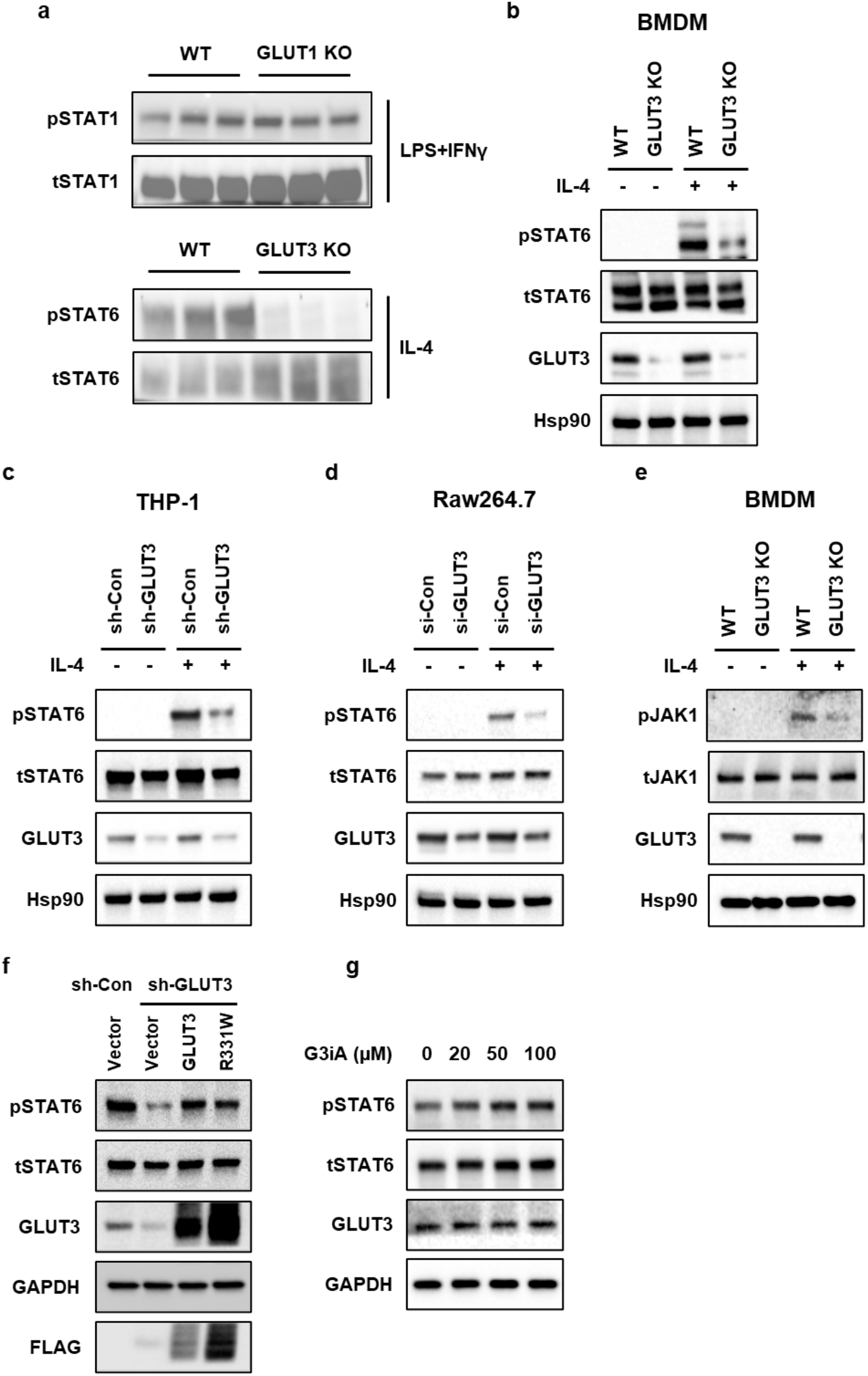
STAT6 signaling is impaired by GLUT3 deficiency. **(a)** Western blot analysis of the expression of phospho-STAT1 (Y701) and total STAT1 in WT and GLUT1 KO BMDMs after LPS and IFNγ stimulation (upper panel). The expression of phospho-STAT6 (Y641) and total STAT6 in WT and GLUT3 KO BMDMs after IL-4 stimulation (lower panel). **(b**) Representative Western blot showing IL-4–mediated protein expression of pSTAT6 and STAT6 in WT and GLUT3 KO BMDMs. **(c-d)** Western blot analysis of the expression of pSTAT6 and STAT6 after knockdown of endogenous GLUT3 in THP-1 cells and RAW 264.7 cells using sh-RNA and si-RNA, respectively. **(e)** The expression of phospho-JAK1 (Y1034/1035) and total JAK1 was analyzed in WT and GLUT3 KO BMDMs after IL-4 stimulation. **(f)** Expression of pSTAT6 and STAT6 after overexpression of wild type GLUT3 or GLUT3 R331W mutant and knockdown of endogenous GLUT3 in THP-1 cells. **(g)** The expression of pSTAT6 and total STAT6 was analyzed in THP-1 cells after treatment with each concentration of G3iA and IL-4.

As GLUT3 knockout did not significantly affect glucose uptake in BMDM, we next investigated whether the glucose transport function of GLUT3 was required for its role in stimulating STAT6 signaling. First, after identifying missense mutations that blocked glucose transport in the GLUT1 transporter without impacting protein stability, the orthologous missense mutations were generated in GLUT3 alleles (Deng *et al.*, 2014; Raja and Kinne, 2020). After lentiviral transduction, WT and most mutant GLUT3 alleles were stably expressed and membrane localized in Rat2 fibroblasts (Supplementary Fig. 1a-b). Among these mutants, the R331W mutant showed the greatest reduction in glucose uptake compared to WT GLUT3 and was selected for downstream analyses (Supplementary Fig. 1c). Short-hairpin resistant alleles of GLUT3 (WT, R331W) were generated and stably expressed in THP-1 cells. Then, sh-GLUT3 was used to knockdown endogenous GLUT3 in THP-1 cells expressing hairpin-resistant WT or R331W mutant GLUT3. As expected, sh-GLUT3 compromised STAT6 signaling in vector control cells. Notably, both WT and R331W GLUT3 rescued pSTAT6 activation after endogenous GLUT3 knockdown (Fig. 4f). The transport-independent role of GLUT3 in STAT6 signal activation was further confirmed through the chemical inhibition of GLUT3. THP-1 cells were treated with a specific, small molecule inhibitor of GLUT3, G3iA (Iancu et al., 2022). After inhibition of GLUT3 with G3iA, the activation of STAT6 by IL-4 was not impaired (Fig. 4g). We next determined whether treatment with G3iA might impact the expression of markers of macrophage polarization. THP-1 cells treated with were treated with M1 or M2 polarization stimuli in the presence of increasing concentrations of G3iA for 24 hrs. Inhibition of GLUT3 transport did not significantly change the expression of either M2 or M1 differentiation markers (Supplementary Fig. 2a, b). We conclude that GLUT3 promotes IL-4/STAT6 signaling and M2 polarization in a glucose transport-independent manner.

### GLUT3 is localized in endosomes

Several previous studies have demonstrated that GLUT3 is largely intracellular rather than localized to the cell surface (Ferreira et al., 2011; McClory et al., 2014). In particular, in primary cortical neurons, GLUT3 is mostly localized in endosomes (McClory *et al.*, 2014). Given previous studies demonstrating that the IL4 receptor complex also undergoes endocytosis and that these endosomes play a role as a signaling platform (Kurgonaite *et al.*, 2015), we next investigated the cellular localization of GLUT3 in macrophages. The localization of GLUT1 and GLUT3 were assessed after fractionation of THP-1 cells. Notably, most GLUT1 was in the plasma membrane fraction, whereas GLUT3 was found predominantly in the intracellular (nonplasma membrane) fraction (Fig. 5a). IF was performed to confirm the biochemical fractionation. Indeed, GLUT1 showed a strong localization on the cell surface, whereas GLUT3 minimally stained the cell surface and instead showed a strong co-localization signal with the endosomal marker early endosomes antigen 1 (EEA1) (Fig. 5b). To determine whether GLUT3 is specifically enriched in the endosomes, we fractionated WT BMDMs and GLUT3 KO BMDMs and enriched for the plasma membrane and endosomes using fractionation kits. Consistent with the immunostaining, GLUT 1 again was found mostly in the plasma membrane fraction, whereas GLUT3 was strongly enriched in the endosomal fraction (Fig. 5c). A similar distribution for GLUT3 was noted in THP-1 and Raw 264.7 cells, where GLUT3 was predominantly present in the endosomes (Fig. 5d-e). In all three cells, the endosomal localization of GLUT3 was independent of IL4. Notably, fractionation also revealed that STAT6 and phospho-STAT6 were also mostly present in the endosomal fraction. Consistent with a critical role for endosome in IL-4 signaling, activation of STAT6 by IL4 was reduced by dynasore (Fig. 5f), a small molecule inhibitor of dynamin and endocytosis.

**Figure 5.**
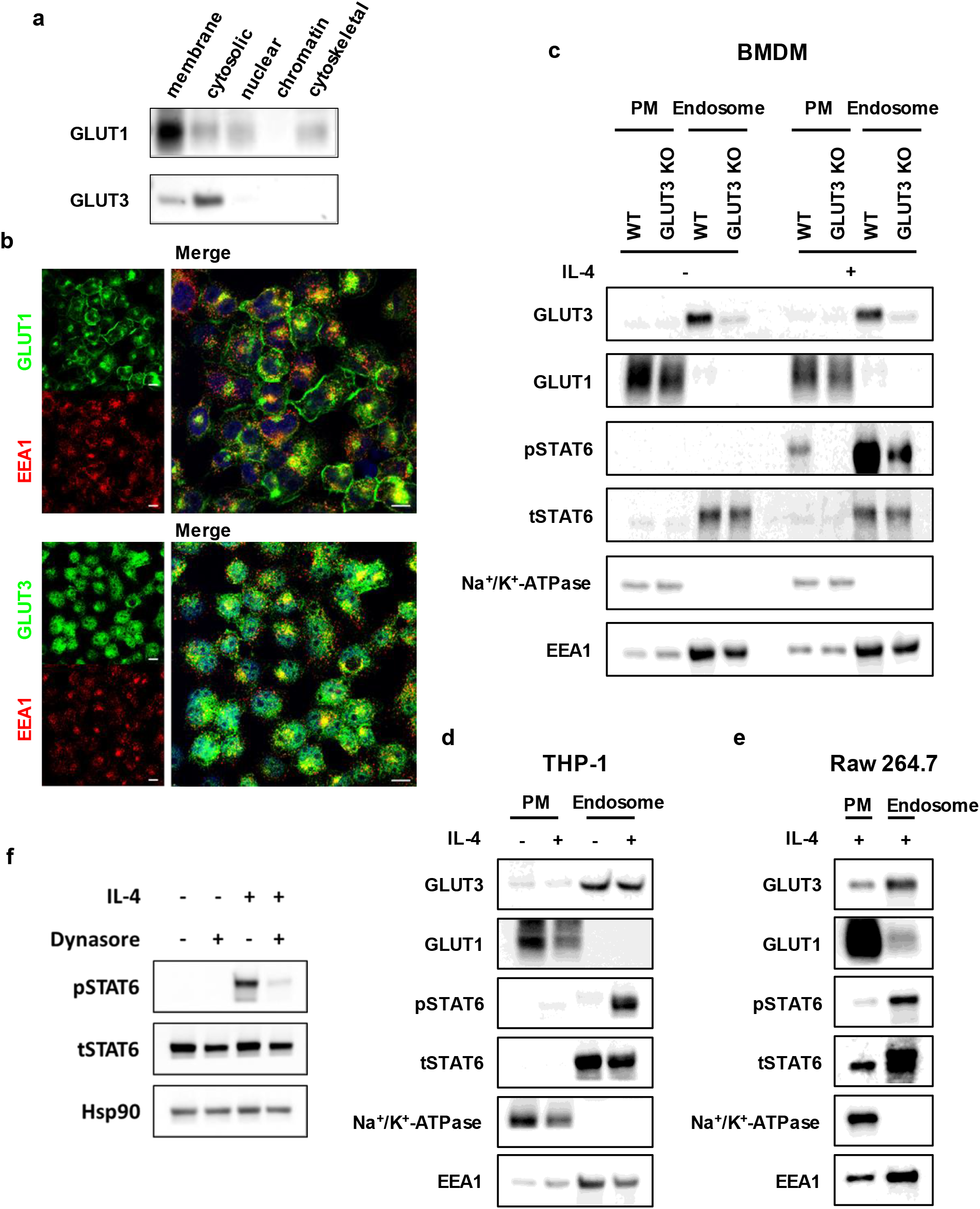
GLUT3 is localized in endosomes. **(a)** Western blot of GLUT1 and GLUT3 in membrane, cytosol, nuclear, chromatin, and cytoskeletal fraction from THP-1 cells. **(b)** Representative immunofluorescence stains of GLUT1 (upper panel, green), GLUT3 (lower panel, green), EEA1 (red), and DAPI (blue) in THP-1 cells. Scale bar = 10 μm. **(c-e)** Western blot analysis of the expression of GLUT3, GLUT1, pSTAT6 and STAT6 in the isolated plasma membrane (PM) and endosome fraction from WT and GLUT3 KO BMDMs (c), THP-1 cells (d) and RAW 264.7 cells (e). Na^+^/K^+^-ATPase and EEA1 were used as fractionation controls for the plasma membrane and endosome, respectively. **(f)** Western blot analysis of the expression of pSTAT6 and STAT6 in THP-1 cells in the presence or absence of IL-4 and dynasore.

### GLUT3 interacts with Ras and regulates IL4R endocytosis

The clathrin-independent endocytosis of IL4R, common gamma chain (gc), and other RTKs is a process that is coordinated by small GTPases, including Rac1 and Ras, and downstream by PAK1/2 and actin (Grassart et al., 2008; Sauvonnet et al., 2005). To dissect how endosomal GLUT3 might regulate IL-4R endocytosis and possibly other endocytic signaling pathways more broadly, we assessed the relationship between GLUT3 and proteins implicated in non-clathrin mediated endocytosis and membrane dynamics. Several proteomic studies have previously demonstrated an interaction between GLUT3 and the Ras GTPases (H-Ras, N-Ras, and K-Ras) (Bigenzahn et al., 2018; Kovalski et al., 2019). We performed a co-immunoprecipitation assay between GLUT3 and Ras, and found that endogenous GLUT3 interacted with Ras in BMDMs (Fig. 6a). To confirm the specificity of the interaction, HEK293T were transfected with GLUT1 or GLUT3. Ras was detected after immunoprecipitation of GLUT3, but not GLUT1, demonstrating the specificity of the interaction between Ras and GLUT3 (Fig. 6b). GLUT1 and GLUT3 are highly homologous proteins, with most differences localizing to their intracytoplasmic loop (ICH) and carboxy terminal (Cterm) motifs (Supplementary Fig. 3). Using GLUT3 chimeric mutants that possessed the GLUT1 ICH, GLUT1 Cterm, or both, Ras co-immunoprecipitation experiments were repeated. Notably, the WT and Cterm GLUT3 alleles interacted with Ras, but not the ICH and double chimeric mutant, indicating that the GLUT3 ICH motif was necessary for the interaction between GLUT3 and Ras (Fig. 6c). Next, short-hairpin resistant WT and chimeric GLUT3 alleles were lentivirally transduced in THP-1 cells. After knockdown of endogenous GLUT3 with sh-GLUT3, pSTAT6 signaling was partially rescued by WT and Cterm GLUT3 alleles, but less effectively rescued by the ICH and double chimeric GLUT3 mutants pSTAT6 activation after endogenous GLUT3 knockdown (Fig. 6d).

**Figure 6.**
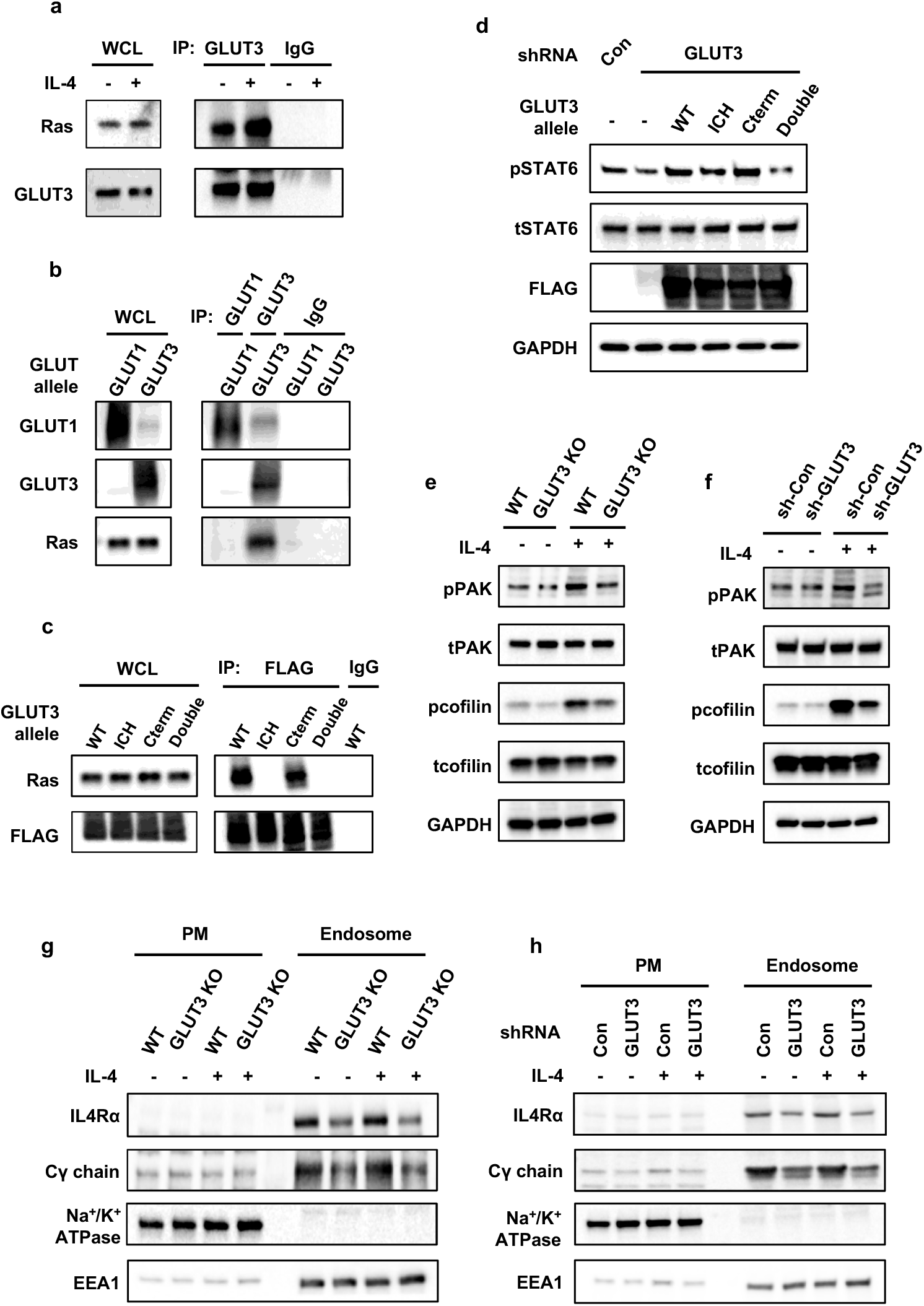
GLUT3 interacts with Ras and regulates endocytosis of the IL4R subunits. **(a)** Interaction between GLUT3 and Ras in WT BMDMs. GLUT3 was immunoprecipitated from the cell lysates and Ras (D2C1 Rabbit mAb recognizing N-Ras and K-Ras) was detected by Western blot. **(b)** HEK293T cells were transfected with the indicated GLUT allele, and GLUT1 (Fisher Scientific, MA1-37783) or GLUT3 (Abcam, ab15311) were immunoprecipitated. Ras was detected by Western blot. Normal mouse IgG for GLUT1 and a normal rabbit IgG for GLUT3 were used as IP controls. **(c)** HEK293T cells were transfected with the indicated GLUT3 allele (see Figure S3) and GLUT3 alleles were Flag immunoprecipitated; Ras was detected by Western blot. IgG indicates a normal mouse IgG as IP control. **(d)** Expression of pSTAT6 and STAT6 after expression of indicated GLUT3 allele and knockdown of endogenous GLUT3 in THP-1 cells. **(e, f)** Protein expression of phospho-PAK, total PAK, phospho-cofilin (S3) and total cofilin in the presence or absence of IL-4 in WT, GLUT3 KO BMDMs (c) and THP-1 cells transfected with sh-control or sh-GLUT3 plasmid (d). **(g, h)** Western blot analysis of the expression of IL4Rα and Cγ chain in the isolated plasma membrane (PM) and endosomal fraction from WT, GLUT3 KO BMDMs (f) and THP-1 cells lentivirally transduced with sh-Con or sh-GLUT3 plasmid (g). Na^+^/K^+^-ATPase and EEA1 were used as fractionation controls.

One established down-stream targets of Ras is Rac1 (Scita et al., 2000), which also regulates the endocytosis of IL-4R in addition to other cytokine receptors (Doherty and McMahon, 2009; Grassart *et al.*, 2008). PAK and cofilin are then targeted by Rac1 to regulate endocytosis (Doherty and McMahon, 2009; Moller et al., 2019). Thus, we checked whether the phosphorylation status of PAK and cofilin might be affected by GLUT3 deficiency. In BMDMs and THP-1 cells, IL-4 treatment induced a notable increase in both phospho-PAK and phospho-cofilin levels. Notably, GLUT3 KO or shRNA inhibited the phosphorylation of both PAK and cofilin (Fig. 6e-f). Finally, endosomal preparations revealed that the endocytosis of IL4Ra and Common γ chain were also reduced by GLUT3 deficiency (Fig. 6g-h). WB and qRT-PCR analyses of IL4Ra and the Cy chain confirmed that their expression was not affected by GLUT3 deficiency (Supplementary Fig. 4). We conclude that GLUT3 interacts with Ras and regulates IL4R endocytosis and signaling and that this requires the ICH domain of GLUT3.

## DISCUSSION

In many cell types, both GLUT1 and GLUT3 are expressed at high levels, yet the specific functions of GLUT3 in cells like macrophages has not been clarified. For the first time, we investigated both the functions of GLUT1 and GLUT3 in myeloid cells. Using myeloid cell-specific GLUT1 and GLUT3 deletion mice, we revealed that the two proteins have strikingly distinct roles in specifying macrophage function. Our studies confirm previous overexpression and myeloid cell (LysM-Cre) GLUT1 deletion studies that have demonstrated an important role for GLUT1 in glucose uptake and M1 macrophage activation(Cho et al., 2020; Freemerman *et al.*, 2019). In contrast, GLUT3 KO macrophages did not show differences in glucose uptake in vitro and showed defects in M2 polarization.

GLUT1 and GLUT3 have previously been reported to possess distinct subcellular localizations in both polarized and non-polarized mammalian cells (Harris et al., 1992; Sakyo and Kitagawa, 2002). Previous studies have indicated that GLUT3 localizes primarily to intracellular, rather than exclusively to the plasma, membrane of neurons (McClory *et al.*, 2014). We extended these findings to macrophages with immunofluorescence and fractionation experiments. We find that GLUT1 localized largely to the plasma membrane, while GLUT3 localized primarily to endosomes. The localization of GLUT3 to intracellular membranes is consistent with its function endosomal signaling and with the absence of a role in glucose uptake from the media. Notably, we observed that GLUT3-positive endosomes function as ‘signaling endosomes’ for IL-4/STAT6 signal transduction. Activation of JAK1, which preferentially occurs at the endosome (Kurgonaite *et al.*, 2015), is inhibited by GLUT3 deficiency. Activated phospho-STAT6 was enriched in the endosomes of WT BMDMs after IL-4 stimulation, but this activation was notably impaired in the endosomes of GLUT3 KO BMDMs. Consistent with the finding that the inhibition of endocytosis with dynasore could inhibit IL-4-STAT6 activation in WT BMDMs, we found that GLUT3 deficiency inhibited IL-4R endocytosis. Thus, in macrophages, we identify an unexpected and essential function for GLUT3 in signal transduction; one that is transport independent.

Our discovery of a critical role for GLUT3 in the coordination of membrane signaling provide context for previous proteomic studies which revealed Ras isoforms (Bigenzahn *et al.*, 2018; Kovalski *et al.*, 2019) and actin (Huttlin et al., 2015) as GLUT3 interacting proteins. We confirmed that GLUT3 interacted with Ras in BMDMs by co-immunoprecipitation and further that this interaction required the intracytoplasmic loop (ICH). The palmitoylation of GLUT1, but not GLUT3, near the ICH motif is necessary for the efficient localization of GLUT 1 to the plasma membrane (Zhang et al., 2021). Additional studies will be necessary to determine whether palmitoylation plays a role in the differential interactions between GLUT1, GLUT3, and Ras. While Ras signaling is thought to occur primarily at the plasma membrane, Ras isoforms also localize to endosomes (Chandra et al., 2011; Tian et al., 2007). Ras and Rac1 participate broadly in endocytosis and membrane remodeling, and we find that GLUT3 deficiency reduced phosphorylation of the Rac1 targets PAK and cofilin in BMDMs. IL-4R subunits are internalized by an actin-dependent endocytosis route (Kurgonaite *et al.*, 2015; Sauvonnet *et al.*, 2005), and our observations of IL-4 activated and GLUT3-dependent changes in phospho-cofilin are consistent with a role for GLUT3 in coordinating these activities. In summary, we propose that GLUT3 is critical for the function of a signaling complex involving Ras, Rac1, Pak, and actin, which ultimately regulate IL-4 receptor mediated signal transduction. Previous studies have suggested the coordinated activation of IL-4 signaling and Rac-Cdc42-Pak activation (Wery-Zennaro et al., 2000) or between JAK1-STAT6 and Ras1-Erk signaling (So et al., 2007). Our model suggests that crosstalk between these pathways could occur through GLUT3 at the level of RTK endocytosis and activation. Additional studies that specifically address the links between IL-4 and other RTKs and the Ras-Rac1-PAK pathway are necessary to confirm and extend our findings.

While we specifically delineated an upstream role for GLUT3 in IL-4-STAT6 signaling in macrophages, it is likely that its role in promoting signal transduction is more broadly conserved. Many RTKs, and cytokine receptors in particular, require endocytosis to endosomes to promote signaling. However, not all cytokines require endocytosis and endosomal enrichment for signal transduction. Specifically, IFN-γ signaling occurs efficiently at the plasma membrane (Blouin and Lamaze, 2013), perhaps explaining why M1 polarization stimuli may not be abrogated by loss of GLUT3. A more detailed catalog of the specific cytokines and signaling pathways that require GLUT3 for optimal signal transduction will require further investigation.

Overall, our findings suggest that GLUT3, in contrast to GLUT1, plays an unexpected role in membrane dynamics and signal transduction. Like M2 macrophages, many of the cell types in which GLUT3 is highly expressed—neurons, melanocytes, Langerhans cells, platelets, and others—share the feature of having extensive, compartmentalized endomembrane systems. We speculate that GLUT3 is particularly important for the regulation and maintenance of these complex endomembrane complexes. Our studies in BMDMs and macrophage lines reveal novel functions for GLUT3 in M2 polarization and signaling. Mouse models of wound healing and atopic dermatitis confirm the critical role of GLUT3 in signaling in vivo and demonstrate the biological importance of this distinct glucose transporter isoforms. It will be interesting to determine whether GLUT3’s role in STAT signaling is conserved in other cell types that express GLUT3.

## METHODS

### Animal studies

This study was performed in strict accordance with the recommendations in the Guide for the Care and Use of Laboratory Animals of the National Institutes of Health. All animal studies were conducted in accordance with institutional guidelines and was approved by the Institutional Animal Care and Use Committee (IACUC), animal protocol number 2015–101166 of the University of Texas Southwestern. All efforts were made to follow the Replacement, Refinement and Reduction guidelines.

Slc2a1^flox/flox^ and Slc2a3 ^flox/flox^ mice were obtained from Dr. E. Dale Abel, and LysM-Cre mice were obtained from The Jackson Laboratory. We generated myeloid cell specific GLUT1 and GLUT3 KO mice by crossing Slc2a1^flox/flox^ mice and Slc2a3^flox/flox^ mice with LysM-Cre mice, respectively.

For calcipotriol (MC903) induced atopic dermatitis model, 1.125 nmol calcipotriol in ethanol was applied to the right ear and the shaved back of mice for 13 days. Wound healing assays were completed as previously described (Zhang *et al.*, 2018). Briefly, an excisional wound was generated on the shaved back skin of mice with a 3mm punch biopsy. Wounds were splinted with a silicone ring and wound healing was measured every two days.

### Preparation of mouse bone-marrow derived macrophages (BMDMs)

Bone marrow was harvested from age-matched male WT (Slc2a1^flox/flox^; LysM-Cre- or Slc2a3^flox/flox^; LysM-Cre-) and myeloid cell-specific GLUT1 KO (Slc2a1^flox/flox^; LysM-Cre+) and GLUT3 KO (Slc2a3^flox/flox^; LysM-Cre+) as previously described with minor modifications (Weischenfeldt and Porse, 2008). BMDMs were generated by culturing marrow cells in poly-L-lysine coated culture plate for 7 days with 50 ng/ml M-CSF in RPMI 1640 (Sigma-Aldrich, R8758) supplemented with 20% FBS, 1X Glutamax (Fisher Scientific, 35050061) and 1X Antibiotic-Antimycotic solution (Thermo Scientific, 15240062). BMDMs were activated using 100 ng/ml LPS and 50 ng/ml IFN-g (for M1) or 10 ng/ml IL-4 (for M2) for 24 h.

### Cell lines and Cell culture

The human macrophage THP-1 cells (ATCC) were culture in RPMI 1640 supplemented with 10% FBS and 1X Antibiotic-Antimycotic solution. THP-1 monocytes are differentiated into macrophages by 36 h incubation with 20 nM phorbol 12-myristate 13-acetate (PMA, Sigma, P8139). The murine macrophage Raw 274.7 cells (ATCC) were cultured in DMEM (Sigma-Aldrich, D5796) supplemented with 10% FBS and 1X Antibiotic-Antimycotic solution. All cells were grown at 37°C with 5% CO_2_.

### [^3^H]2-deoxyglucose uptake assay

2-DG uptake were measured as previously described (Lee et al., 2015). Briefly, BMDMs from WT, GLUT1 KO and GLUT3 KO mice were seeded in triplicate into 12-well plates overnight. The cells were washed twice with PBS, incubated in basic serum-free DMEM medium for 2 h per well. Uptake was initiated by addition of 1 μCi [3H]2-DG (25-30 Ci/mmol, PerkinElmer, NET549) and 0.1 mM unlabeled 2-DG (Sigma, D8375) to each well for 10 mins. Transport activity was terminated by rapid removal of the uptake medium and subsequent washing three times with cold PBS with 25 mM glucose (Sigma, G7528). Cells were lysed with 0.5 ml of 0.5 M NaOH (Fisher Scientific, SS255-1) and neutralized with 0.5 ml or 0.5 M HCl (Sigma, 320331), which was added and mixed well. 250 μl of the lysate was transferred to a scintillation vial containing scintillation solution, and the sample was analyzed by liquid scintillation counting. Protein concentrations were determined through BCA assays (Thermo Scientific, 23227).

### GLUT3 short hairpins, GLUT3 expression alleles, and lentiviral transductions

GLUT3 short hairpin RNA (shRNA) sequences were designed for pLKO.1 using the TRC shRNA Design Process (https://portals.broadinstitute.org/gpp/public/resources/rules). Forward and reverse oligos were annealed in NEB buffer 2, boiled at 95 for 10 min and slowly cool to room temperature, then ligated into pLKO.1 vector (Addgene #10878) using AgeI/EcoRI. The constructs were confirmed by Sanger Sequencing. An amino-terminal 3xFlag epitope tagged human GLUT3 was generated by PCR (Addgene #72877) (Supplementary Table 1). Missense mutant (N32S, G312S, N315T, R331W) and sh-GLUT3 resistant GLUT3 alleles (sh1sh3 resist) were designed and synthesized as DNA fragments by Integrated DNA Technology (IDT) (Supplementary Table 1). After PCR amplification, GLUT3 missense mutants, the GLUT3 sh1sh3 resistant mutant, and double mutants were cloned by restriction digestion. Short hairpin resistant chimeric mutants of Flag-tagged GLUT3 alleles were generated using NEBuilder HiFi DNA Assembly Cloning Kit (New England Biolabs, E5520) according to the manufacturer’s protocol. Vector (WT GLUT3) and inserts (GLUT3 ICH, GLUT3 Cterm) were amplified by PCR using indicated the primer sets (Supplementary Table 1). PCR fragments were assembled after incubation at 50 °C for 30 min with the NEBuilder^®^ HiFi DNA Assembly Master Mix in the kit. All constructs were confirmed by Sanger sequencing.

For generating lentiviruses, LentiX-293T cells (Clontech, 632180) were seeded at ~60% confluence in antibiotics free media 12–16 h before transfection. 4.5μg of shRNA or expression plasmid, 2.5μg of pMD2.G (Addgene #12259) and 4.5μg of psPAX2 (Addgene #12260) were co-transfected into LentiX-293T cells using Lipofectamine 3000 (Thermo Scientific, L3000015) according to the manufacturer’s protocol. Viruses were collected after 48 and 72 h transfection. For lentiviral transduction, THP-1 cells were seeded into 6-well plate at 70% confluence. Viral supernatant was then added to the cell with the polybrene at a concentration of 8 μg/mL. Cells were selected with puromycin antibiotic at a concentration of 2 μg/mL and hygromycin at a concentration of 100ug/mL. For double transductions (GLUT3 allele + sh-GLUT3), unmodified THP-1 cells were serially transduced first with the GLUT3 expression plasmid by puromycin selection followed by the sh-GLUT3 allele by hygromycin selection.

### siRNA interference

siRNA targeting mouse Slc2a3 was synthesized from Sigma-Aldrich. Cells were transfected with 100 nM siRNA using Lipofectamine RNAiMAX reagent (Thermo Fisher Scientific, 13778150) according to the manufacturer’s protocol.

### RNA extraction and qRT–PCR

RNA was extracted form cells or tissue using a RNeasy Mini Kit (Qiagen, 74106) and reverse transcribed to cDNAs using Iscript cDNA Synthesis Kit (Bio-Rad, 1708891) according to the manufacturer’s protocol. qRT-PCR analyses were performed using the cDNAs from the reverse transcription reactions, gene-specific primers and PowerUp SYBR Green (Applied Biosystems, A25779). All primers for qRT-PCR are listed in Table S1 and S2.

### Isolation of plasma membrane and endosome

Cells were fractionated into plasma membrane and endosome using fractionation kits (Invent Biotechnologies, SM-005 and ED-028) according to the manufacturer’s protocol. Briefly, cells were lysed with the supplied buffer and intact nuclei and un-ruptured cells were removed by the filter cartridge and brief centrifugation. The supernatant was incubated with the supplied precipitation buffer to isolate and enrich the plasma membrane or endosome.

### Immunoblotting and immunoprecipitation

For STAT6 signaling experiments, cells were stimulated with 20ng/ml IL-4 for 30min. 200μM of dynasore (Sigma, D7693) was pretreated for 30 minutes before IL-4 treatment. G3iA was obtained from Dr. Jun-yong Choe and cells pretreated for 10 minutes before IL-4 treatment. After stimulation, cells were lysed with Cell Lysis Buffer (Cell Signaling Technology, 9803) and whole-cell lysates (WCL) were subjected to Laemmli Sample Buffer (Bio-Rad, 1610747) and sodium dodecyl sulfate–polyacrylamide gel electrophoresis (SDS–PAGE). The separated proteins were transferred to a nitrocellulose membrane and incubated with GLUT3 (Abcam, ab191071 or ab15311), GLUT1 (Millipore Sigma, 07-1401), phospho-STAT1 (Y701) (Cell Signaling Technology, 7649), STAT1 (Cell Signaling Technology, 14994), phospho-STAT6 (Y641) (Cell Signaling Technology, 9361), STAT6 (Cell Signaling Technology, 5397), phospho-JAK1(Y1034/1035) (Cell Signaling Technology, 3331), JAK1 (Cell Signaling Technology, 3332), Na^+^/K^+^-ATPase (Cell Signaling Technology, 3010), EEA1 (Cell Signaling Technology, 48453), phospho-PAK (PAK1(T423)/PAK2(T402) (Cell Signaling Technology, 2601), PAK1 (Cell Signaling Technology, 2602), and Ras (Cell Signaling Technology, 8955, D2C1), and Hsp90 (Cell Signaling Technology, 4877) primary antibodies. After incubating with horseradish peroxidase (HRP)-conjugated secondary antibodies, antigens were visualized using a Western Lightning Plus-ECL (Perkin Elmer, 50-904-9323).

For immunoprecipitation, cells were lysed in buffer containing 20 mM Tris–HCl (pH 7.4), 137 mM NaCl, 1 mM MgCl2, 1 mM CaCl2. WCL were incubated with GLUT3 antibody overnight and then with Protein A/G PLUS-Agarose bead slurry (Santa Cruz Biotechnology, sc-2003) for 3h. Immunoprecipitates were analyzed by immunoblotting.

### Immunofluorescence staining

For immunofluorescence staining of cells, cells were fixed with 4% paraformaldehyde in PBS for 10 min, permeabilized with 0.1% Triton X-100 for 10 min and blocked with blocking solution (3% BSA in PBS) for 1 h. Cells were then incubated with GLUT1 (Millipore Sigma, 07-1401), GLUT3 (Santacruz Biotechnology, sc-30107) and EEA1 (Cell Signaling Technology, 48453) primary antibodies in blocking buffer overnight at 4°C, followed by 1 h of incubation with fluorescent dye-labeled secondary antibodies (Thermo Scientific, A11005 or A11008). After mounting with Cytoseal 60 (Thermo Scientific, 23244257), confocal images were captured on an LSM 780 confocal microscope (Zeiss).

For immunofluorescence staining of tissues, tissues were fixed in 4% paraformaldehyde and embedded in paraffin. Sections (5 μm) were deparaffinized, heat retrieved at 95 °C for 30min with citrate buffer (Thermo Scientific, AP-9003-125), permeabilized with 0.1% Triton X-100 for 10 min and blocked with 5% goat-serum and 0.2% BSA in PBS. Tissues were then incubated with Arg1 (Cell Signaling Technology, 93668) and F4/80 (Thermo Scientific, MA1-91124) primary antibodies overnight at 4°C, followed by 1 h of incubation with fluorescent dye-labeled secondary antibodies (Thermo Scientific, A11005 or A11007). After mounting with ProLong Gold antifade reagent with DAPI (Thermo Scientific, P36935), confocal images were captured on an LSM 780 confocal microscope (Zeiss).

---

**Supplementary Figure 1.**
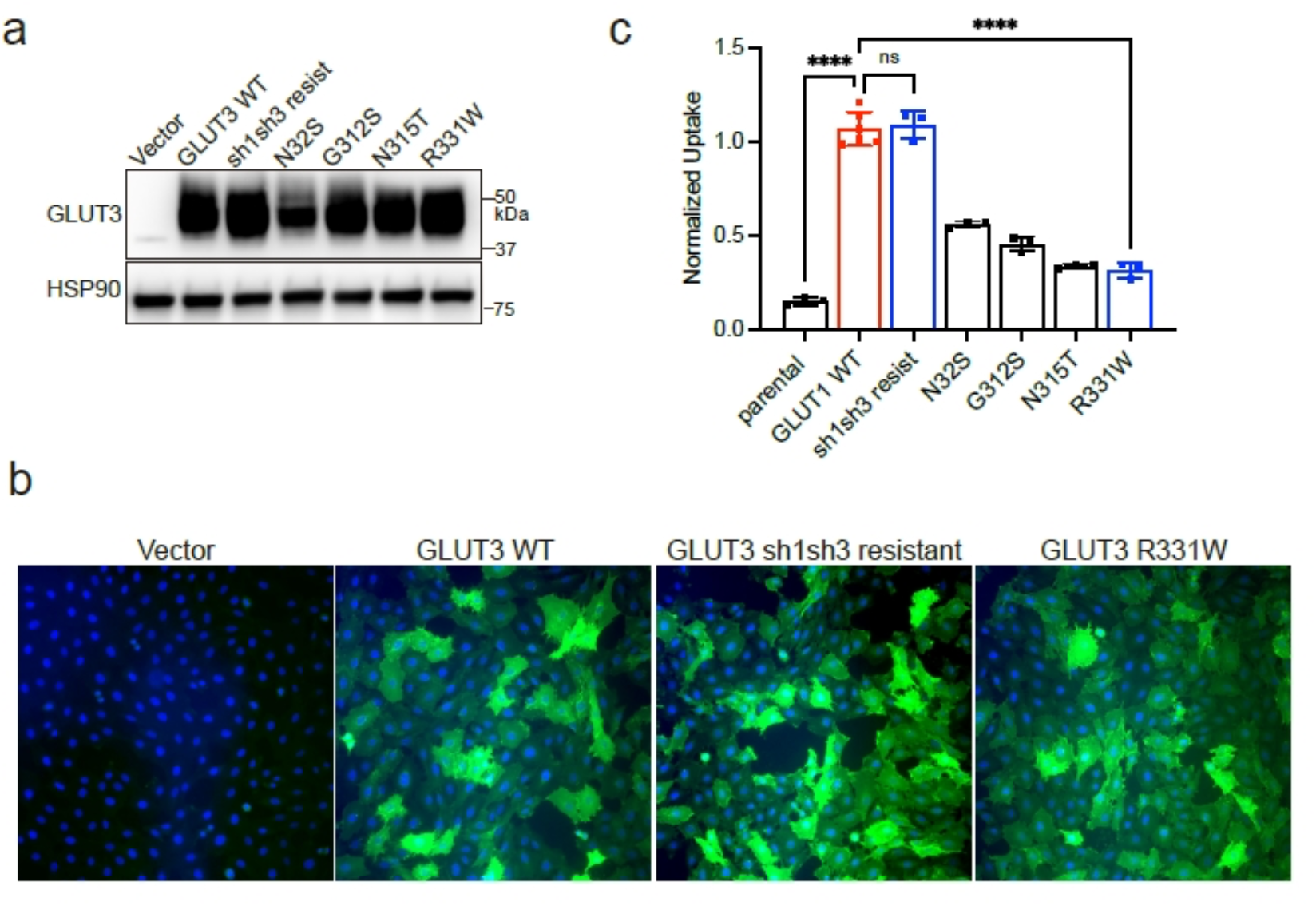
Expression and glucose uptake of WT GLUT3 and GLUT3 mutants in Rat2 fibroblasts. **(a)** Western blot analysis of GLUT3 mutant alleles in Rat2 fibroblasts after lentiviral transduction of the indicated plasmid. HSP90, loading control. **(b)** Representative immunofluorescence stains of Flag-GLUT3 (green) and DAPI (blue) in vector, GLUT3 WT, GLUT3 sh1sh3 resistant, and GLUT3 R331W expressing Rat2 fibroblasts. (c) 2-Deoxy-D-glucose uptake in Rat2 fibroblasts expressing each plasmid. Data shown as mean ± SEM. P values were calculated by one-way ANOVA with Dunnett tests. ****P < 0.0001.

**Supplementary Figure 2.**
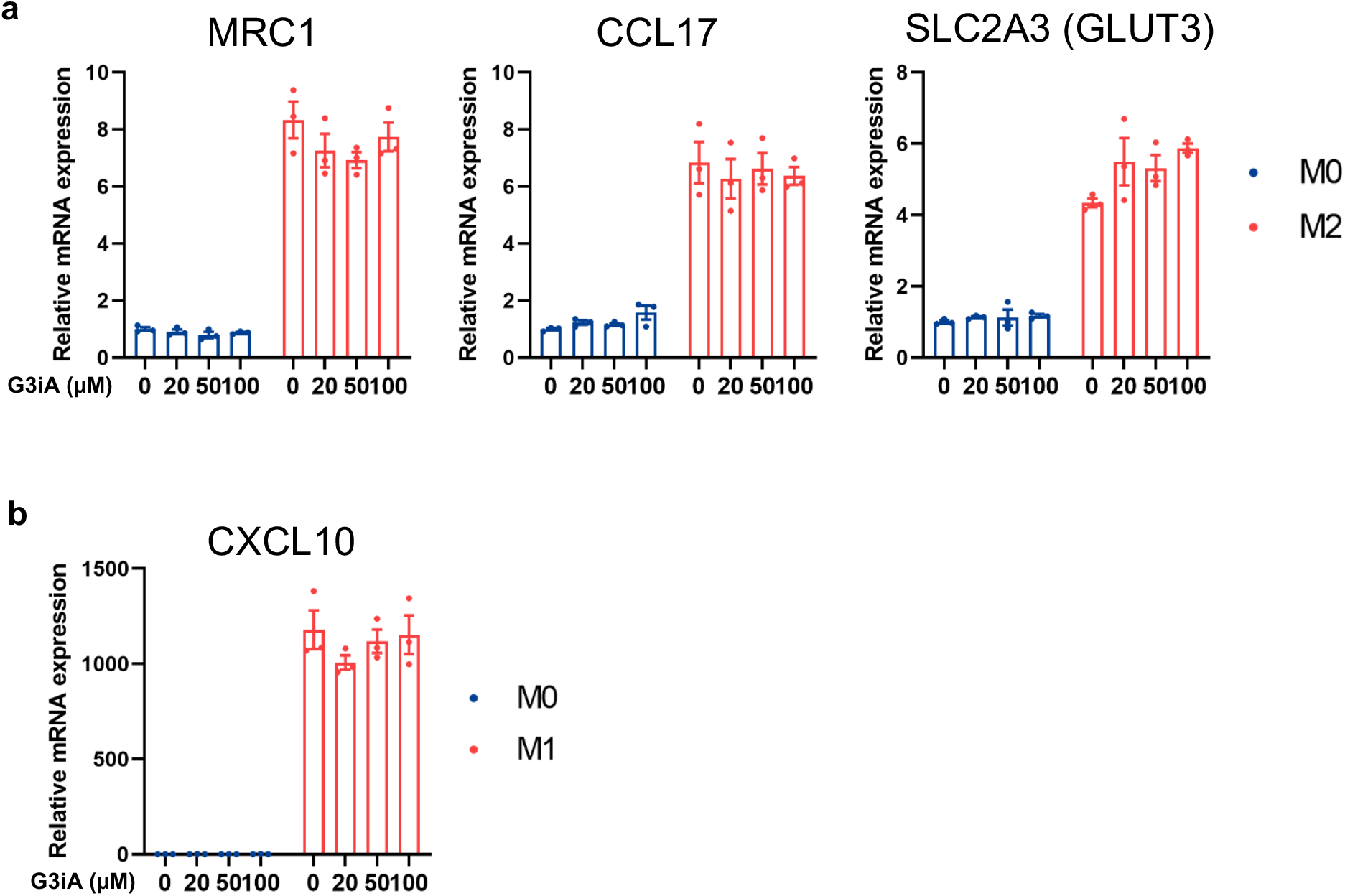
Expression of macrophage polarization markers in THP-1 cells after treatment with chemical inhibitor of GLUT3. **(a)** THP-1 cells were induced with IL-4 and the indicated concentration of G3iA for 24 hours. Then mRNA expression of the indicated M2 polarization marker was assessed. **(b)** THP-1 cells were induced with LPS + IFN-γ and the indicated concentration of G3iA for 24 hours. Then mRNA expression of CXCL10 (M1 polarization marker) was assessed.

**Supplementary Figure 3.**
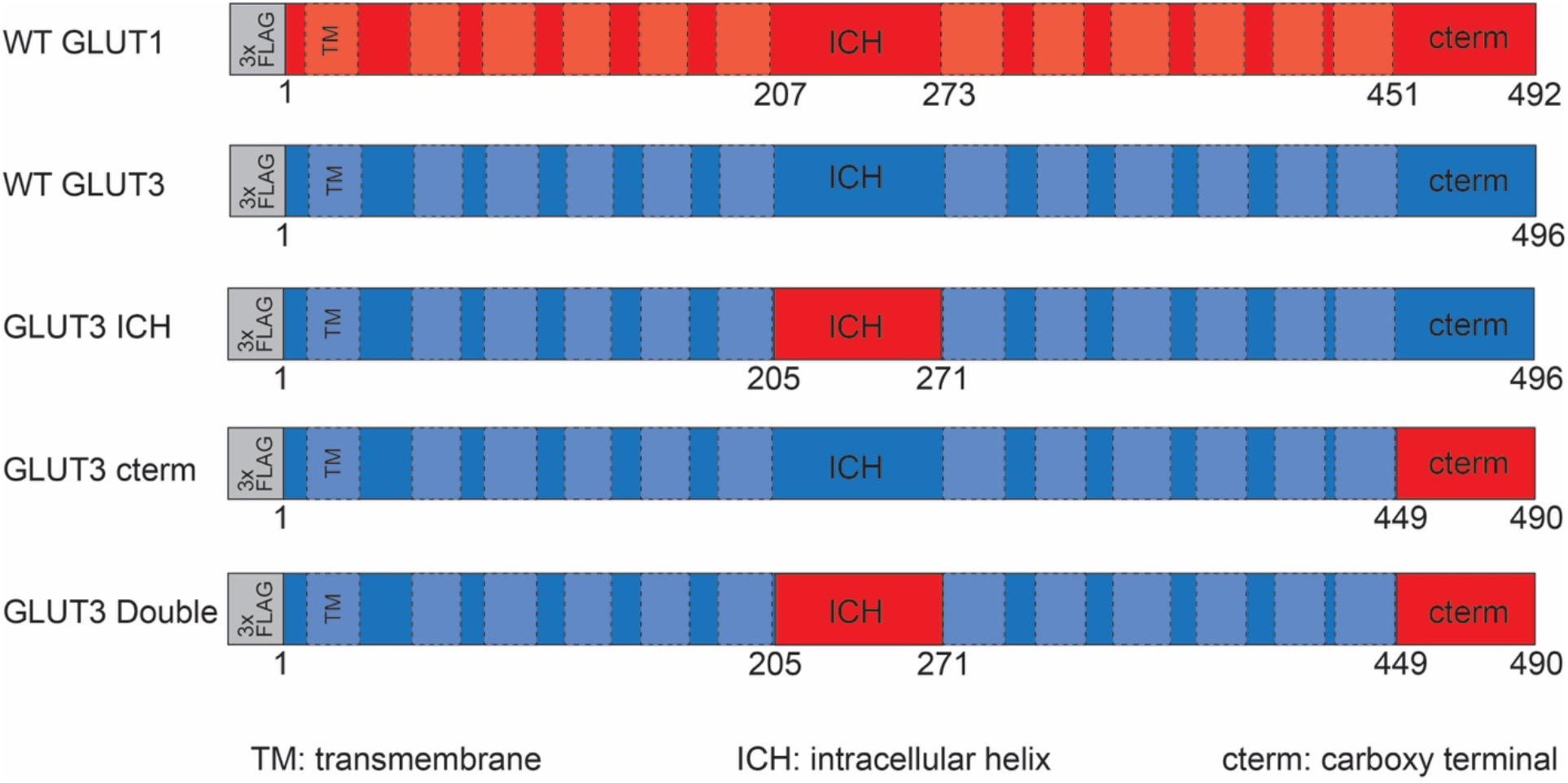
Schematic of chimeric GLUT1/GLUT3 mutants used in Ras coimmunoprecipitation experiments. Alleles were amino-terminally tagged with a 3xFlag epitope tag. Dashed boxes indicated predicted transmembrane (TM) domains.

**Supplementary Figure 4.**
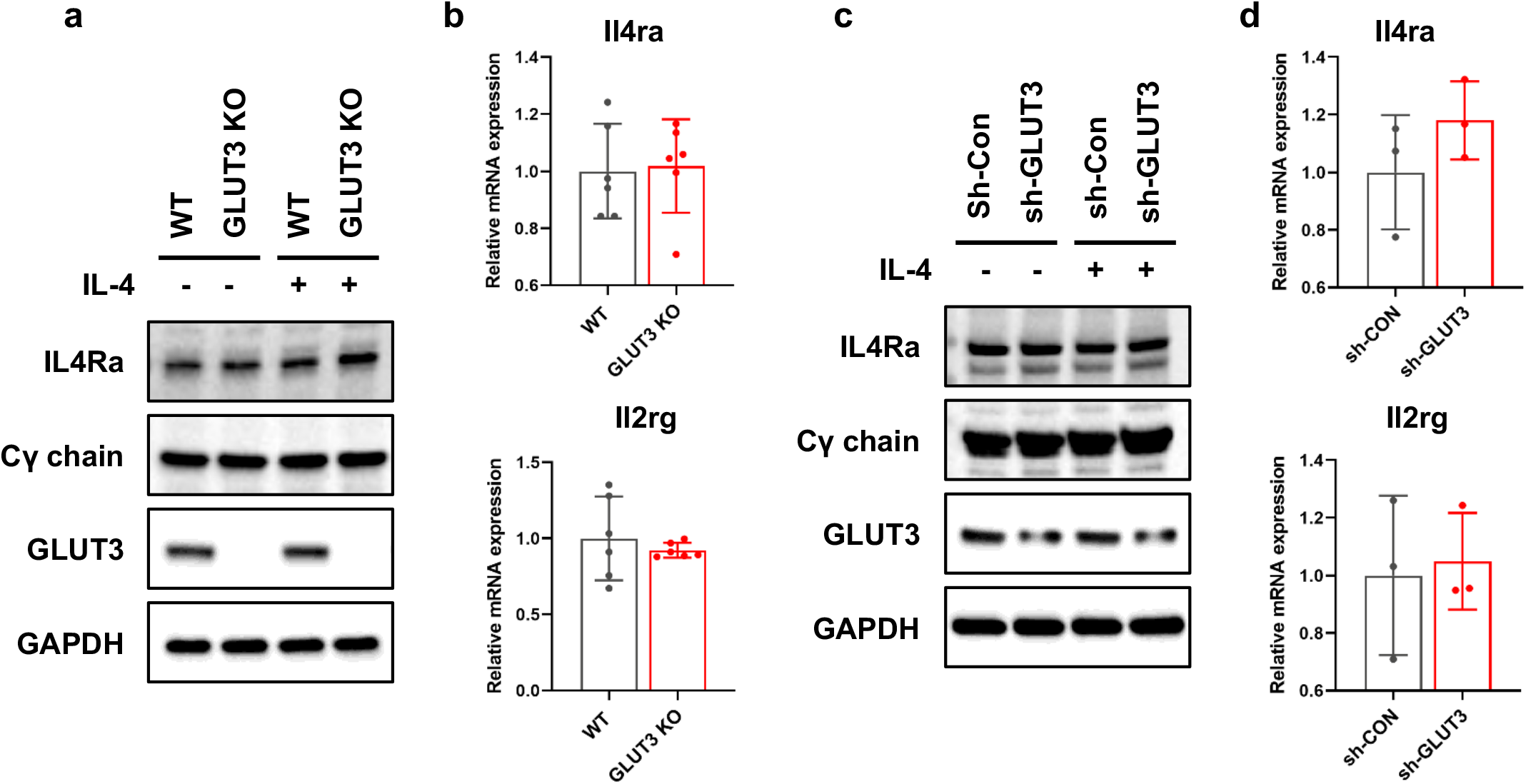
Expression of membrane receptors in GLUT3 KO BMDM and THP-1 cells after GLUT3 shRNA. **(a)** Western blot for IL4Rα and Cγ chain in WT and GLUT3 KO BMDMs. GAPDH, loading control. **(b)** mRNA levels of IL4Rα and Cγ chain transcripts in WT and GLUT3 KO BMDMs. **(c)** Western blot for IL4Rα and Cγ chain in WT and GLUT3 KO IL4Rα and Cγ chain in THP-1 cells transduced with sh-Con or sh-GLUT3 plasmid. GAPDH, loading control. **(d)** mRNA levels of IL4Rα and Cγ chain transcripts in THP-1 cells transduced with sh-Con or sh-GLUT3 plasmid.

**Supplementary Figure 5.**
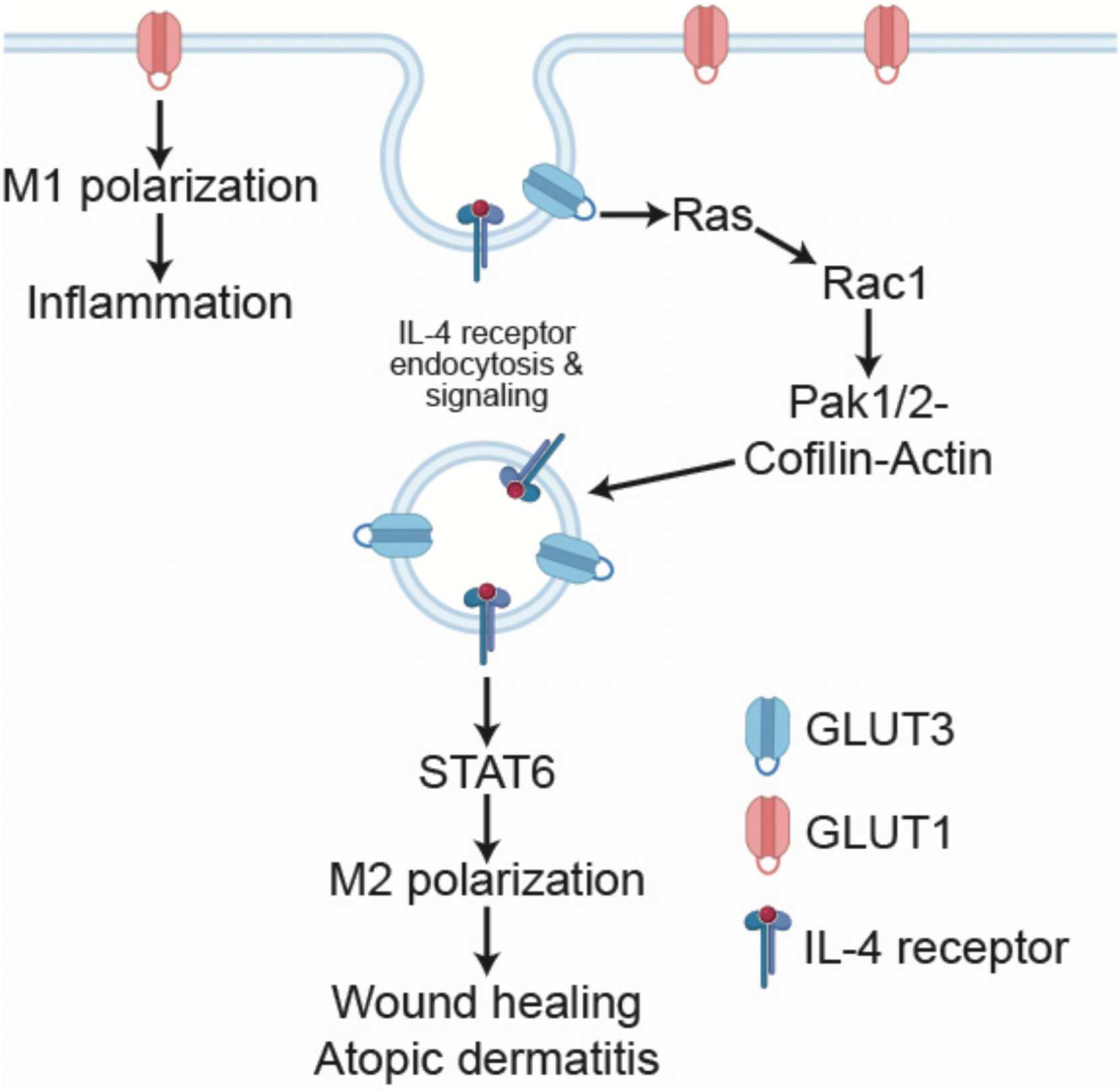
Summary graphic

## Notes

### Competing Interest Statement

The authors have declared no competing interest.

## References

Bigenzahn, J.W., Collu, G.M., Kartnig, F., Pieraks, M., Vladimer, G.I., Heinz, L.X., Sedlyarov, V., Schischlik, F., Fauster, A., Rebsamen, M., et al. (2018). LZTR1 is a regulator of RAS ubiquitination and signaling. Science 362, 1171–1177. 10.1126/science.aap8210.

Blouin, C.M., and Lamaze, C. (2013). Interferon gamma receptor: the beginning of the journey. Front Immunol 4, 267. 10.3389/fimmu.2013.00267.

Chandra, A., Grecco, H.E., Pisupati, V., Perera, D., Cassidy, L., Skoulidis, F., Ismail, S.A., Hedberg, C., Hanzal-Bayer, M., Venkitaraman, A.R., et al. (2011). The GDI-like solubilizing factor PDEdelta sustains the spatial organization and signalling of Ras family proteins. Nat Cell Biol 14, 148–158. 10.1038/ncb2394.

Cho, H., Kwon, H.Y., Sharma, A., Lee, S.H., Liu, X., Miyamoto, N., Kim, J.J., Im, S.H., Kang, N.Y., and Chang, Y.T. (2022). Visualizing inflammation with an M1 macrophage selective probe via GLUT1 as the gating target. Nat Commun 13, 5974. 10.1038/s41467-022-33526-z.

Cho, S.J., Moon, J.S., Nikahira, K., Yun, H.S., Harris, R., Hong, K.S., Huang, H., Choi, A.M.K., and Stout-Delgado, H. (2020). GLUT1-dependent glycolysis regulates exacerbation of fibrosis via AIM2 inflammasome activation. Thorax 75, 227–236. 10.1136/thoraxjnl-2019-213571.

Cura, A.J., and Carruthers, A. (2012). Role of monosaccharide transport proteins in carbohydrate assimilation, distribution, metabolism, and homeostasis. Comprehensive Physiology 2, 863–914. 10.1002/cphy.c110024.

Deng, D., Sun, P., Yan, C., Ke, M., Jiang, X., Xiong, L., Ren, W., Hirata, K., Yamamoto, M., Fan, S., and Yan, N. (2015). Molecular basis of ligand recognition and transport by glucose transporters. Nature 526, 391396. 10.1038/nature14655.

Deng, D., Xu, C., Sun, P., Wu, J., Yan, C., Hu, M., and Yan, N. (2014). Crystal structure of the human glucose transporter GLUT1. Nature 510, 121–125. 10.1038/nature13306.

Doherty, G.J., and McMahon, H.T. (2009). Mechanisms of endocytosis. Annu Rev Biochem 78, 857–902. 10.1146/annurev.biochem.78.081307.110540.

Ferreira, J.M., Burnett, A.L., and Rameau, G.A. (2011). Activity-dependent regulation of surface glucose transporter-3. J Neurosci 31, 1991–1999. 10.1523/JNEUROSCI.1850-09.2011.

Fidler, T.P., Campbell, R.A., Funari, T., Dunne, N., Balderas Angeles, E., Middleton, E.A., Chaudhuri, D., Weyrich, A.S., and Abel, E.D. (2017). Deletion of GLUT1 and GLUT3 Reveals Multiple Roles for Glucose Metabolism in Platelet and Megakaryocyte Function. Cell Rep 20, 881–894. 10.1016/j.celrep.2017.06.083.

Freemerman, A.J., Zhao, L., Pingili, A.K., Teng, B., Cozzo, A.J., Fuller, A.M., Johnson, A.R., Milner, J.J., Lim, M. F., Galanko, J.A., et al. (2019). Myeloid Slc2a1-Deficient Murine Model Revealed Macrophage Activation and Metabolic Phenotype Are Fueled by GLUT1. J Immunol 202, 1265–1286. 10.4049/jimmunol.1800002.

Fu, Y., Maianu, L., Melbert, B.R., and Garvey, W.T. (2004). Facilitative glucose transporter gene expression in human lymphocytes, monocytes, and macrophages: a role for GLUT isoforms 1, 3, and 5 in the immune response and foam cell formation. Blood Cells Mol Dis 32, 182–190. 10.1016/j.bcmd.2003.09.002.

Grassart, A., Dujeancourt, A., Lazarow, P.B., Dautry-Varsat, A., and Sauvonnet, N. (2008). Clathrin-independent endocytosis used by the IL-2 receptor is regulated by Rac1, Pak1 and Pak2. EMBO Rep 9, 356–362. 10.1038/embor.2008.28.

Grimes, M.L., Zhou, J., Beattie, E.C., Yuen, E.C., Hall, D.E., Valletta, J.S., Topp, K.S., LaVail, J.H., Bunnett, N. W., and Mobley, W.C. (1996). Endocytosis of activated TrkA: evidence that nerve growth factor induces formation of signaling endosomes. J Neurosci 16, 7950–7964.

Harris, D.S., Slot, J.W., Geuze, H.J., and James, D.E. (1992). Polarized distribution of glucose transporter isoforms in Caco-2 cells. Proc Natl Acad Sci U S A 89, 7556–7560. 10.1073/pnas.89.16.7556.

Hu, X., Li, J., Fu, M., Zhao, X., and Wang, W. (2021). The JAK/STAT signaling pathway: from bench to clinic. Signal Transduct Target Ther 6, 402. 10.1038/s41392-021-00791-1.

Huttlin, E.L., Ting, L., Bruckner, R.J., Gebreab, F., Gygi, M.P., Szpyt, J., Tam, S., Zarraga, G., Colby, G., Baltier, K., et al. (2015). The BioPlex Network: A Systematic Exploration of the Human Interactome. Cell 162, 425–440. 10.1016/j.cell.2015.06.043.

Iancu, C.V., Bocci, G., Ishtikhar, M., Khamrai, M., Oreb, M., Oprea, T.I., and Choe, J.Y. (2022). GLUT3 inhibitor discovery through in silico ligand screening and in vivo validation in eukaryotic expression systems. Sci Rep 12, 1429. 10.1038/s41598-022-05383-9.

Kasraie, S., and Werfel, T. (2013). Role of macrophages in the pathogenesis of atopic dermatitis. Mediators Inflamm 2013, 942375. 10.1155/2013/942375.

Koo, J.H., Jang, H.Y., Lee, Y., Moon, Y.J., Bae, E.J., Yun, S.K., and Park, B.H. (2019). Myeloid cell-specific sirtuin 6 deficiency delays wound healing in mice by modulating inflammation and macrophage phenotypes. Exp Mol Med 51, 1–10. 10.1038/s12276-019-0248-9.

Kovalski, J.R., Bhaduri, A., Zehnder, A.M., Neela, P.H., Che, Y., Wozniak, G.G., and Khavari, P.A. (2019). The Functional Proximal Proteome of Oncogenic Ras Includes mTORC2. Mol Cell 73, 830–844 e812. 10.1016/j.molcel.2018.12.001.

Kurgonaite, K., Gandhi, H., Kurth, T., Pautot, S., Schwille, P., Weidemann, T., and Bokel, C. (2015). Essential role of endocytosis for interleukin-4-receptor-mediated JAK/STAT signalling. J Cell Sci 128, 3781–3795. 10.1242/jcs.170969.

Lee, E.E., Ma, J., Sacharidou, A., Mi, W., Salato, V.K., Nguyen, N., Jiang, Y., Pascual, J.M., North, P.E., Shaul, P.W., et al. (2015). A Protein Kinase C Phosphorylation Motif in GLUT1 Affects Glucose Transport and is Mutated in GLUT1 Deficiency Syndrome. Mol Cell 58, 845–853. 10.1016/j.molcel.2015.04.015.

Ley, K. (2017). M1 Means Kill; M2 Means Heal. J Immunol 199, 2191–2193. 10.4049/jimmunol.1701135.

Li, M., Hener, P., Zhang, Z., Kato, S., Metzger, D., and Chambon, P. (2006). Topical vitamin D3 and low-calcemic analogs induce thymic stromal lymphopoietin in mouse keratinocytes and trigger an atopic dermatitis. Proc Natl Acad Sci U S A 103, 11736–11741. 10.1073/pnas.0604575103.

Li, M., Messaddeq, N., Teletin, M., Pasquali, J.L., Metzger, D., and Chambon, P. (2005). Retinoid X receptor ablation in adult mouse keratinocytes generates an atopic dermatitis triggered by thymic stromal lymphopoietin. Proc Natl Acad Sci U S A 102, 14795–14800. 10.1073/pnas.0507385102.

Mantovani, A., Sica, A., and Locati, M. (2005). Macrophage polarization comes of age. Immunity 23, 344346. 10.1016/j.immuni.2005.10.001.

McClory, H., Williams, D., Sapp, E., Gatune, L.W., Wang, P., DiFiglia, M., and Li, X. (2014). Glucose transporter 3 is a rab11-dependent trafficking cargo and its transport to the cell surface is reduced in neurons of CAG140 Huntington’s disease mice. Acta Neuropathol Commun 2, 179. 10.1186/s40478-014-0178-7.

Moller, L.L.V., Klip, A., and Sylow, L. (2019). Rho GTPases-Emerging Regulators of Glucose Homeostasis and Metabolic Health. Cells 8. 10.3390/cells8050434.

Munoz-Rojas, A.R., Kelsey, I., Pappalardo, J.L., Chen, M., and Miller-Jensen, K. (2021). Co-stimulation with opposing macrophage polarization cues leads to orthogonal secretion programs in individual cells. Nat Commun 12, 301. 10.1038/s41467-020-20540-2.

Murray, P.J. (2017). Macrophage Polarization. Annu Rev Physiol 79, 541–566. 10.1146/annurev-physiol-022516-034339.

Navale, A.M., and Paranjape, A.N. (2016). Glucose transporters: physiological and pathological roles. Biophys Rev 8, 5–9. 10.1007/s12551-015-0186-2.

Oetjen, L.K., Mack, M.R., Feng, J., Whelan, T.M., Niu, H., Guo, C.J., Chen, S., Trier, A.M., Xu, A.Z., Tripathi, S.V., et al. (2017). Sensory Neurons Co-opt Classical Immune Signaling Pathways to Mediate Chronic Itch. Cell 171, 217–228 e213. 10.1016/j.cell.2017.08.006.

Okonkwo, U.A., Chen, L., Ma, D., Haywood, V.A., Barakat, M., Urao, N., and DiPietro, L.A. (2020). Compromised angiogenesis and vascular Integrity in impaired diabetic wound healing. PLoS One 15, e0231962. 10.1371/journal.pone.0231962.

Orecchioni, M., Ghosheh, Y., Pramod, A.B., and Ley, K. (2019). Macrophage Polarization: Different Gene Signatures in M1(LPS+) vs. Classically and M2(LPS-) vs. Alternatively Activated Macrophages. Front Immunol 10, 1084. 10.3389/fimmu.2019.01084.

Raja, M., and Kinne, R.K.H. (2020). Mechanistic Insights into Protein Stability and Self-aggregation in GLUT1 Genetic Variants Causing GLUT1-Deficiency Syndrome. J Membr Biol 253, 87–99. 10.1007/s00232-020-00108-3.

Rehak, L., Giurato, L., Meloni, M., Panunzi, A., Manti, G.M., and Uccioli, L. (2022). The Immune-Centric Revolution in the Diabetic Foot: Monocytes and Lymphocytes Role in Wound Healing and Tissue Regeneration-A Narrative Review. J Clin Med 11. 10.3390/jcm11030889.

Sakyo, T., and Kitagawa, T. (2002). Differential localization of glucose transporter isoforms in non-polarized mammalian cells: distribution of GLUT1 but not GLUT3 to detergent-resistant membrane domains. Biochim Biophys Acta 1567, 165–175. 10.1016/s0005-2736(02)00613-2.

Sauvonnet, N., Dujeancourt, A., and Dautry-Varsat, A. (2005). Cortactin and dynamin are required for the clathrin-independent endocytosis of gammac cytokine receptor. J Cell Biol 168, 155–163. 10.1083/jcb.200406174.

Scita, G., Tenca, P., Frittoli, E., Tocchetti, A., Innocenti, M., Giardina, G., and Di Fiore, P.P. (2000). Signaling from Ras to Rac and beyond: not just a matter of GEFs. EMBO J 19, 2393–2398. 10.1093/emboj/19.11.2393.

Sica, A., and Mantovani, A. (2012). Macrophage plasticity and polarization: in vivo veritas. J Clin Invest 122, 787–795. 10.1172/JCI59643.

Simpson, I.A., Dwyer, D., Malide, D., Moley, K.H., Travis, A., and Vannucci, S.J. (2008). The facilitative glucose transporter GLUT3: 20 years of distinction. Am J Physiol Endocrinol Metab 295, E242–253. 10.1152/ajpendo.90388.2008.

So, E.Y., Oh, J., Jang, J.Y., Kim, J.H., and Lee, C.E. (2007). Ras/Erk pathway positively regulates Jak1/STAT6 activity and IL-4 gene expression in Jurkat T cells. Mol Immunol 44, 3416–3426. 10.1016/j.molimm.2007.02.022.

Sorkin, A., and von Zastrow, M. (2009). Endocytosis and signalling: intertwining molecular networks. Nat Rev Mol Cell Biol 10, 609–622. 10.1038/nrm2748.

Suzuki, K., Meguro, K., Nakagomi, D., and Nakajima, H. (2017). Roles of alternatively activated M2 macrophages in allergic contact dermatitis. Allergol Int 66, 392–397. 10.1016/j.alit.2017.02.015.

Tian, T., Harding, A., Inder, K., Plowman, S., Parton, R.G., and Hancock, J.F. (2007). Plasma membrane nanoswitches generate high-fidelity Ras signal transduction. Nat Cell Biol 9, 905–914. 10.1038/ncb1615.

Vieira, A.V., Lamaze, C., and Schmid, S.L. (1996). Control of EGF receptor signaling by clathrin-mediated endocytosis. Science 274, 2086–2089. 10.1126/science.274.5295.2086.

Weischenfeldt, J., and Porse, B. (2008). Bone Marrow-Derived Macrophages (BMM): Isolation and Applications. CSH Protoc 2008, pdb prot5080. 10.1101/pdb.prot5080.

Wery-Zennaro, S., Zugaza, J.L., Letourneur, M., Bertoglio, J., and Pierre, J. (2000). IL-4 regulation of IL-6 production involves Rac/Cdc42-and p38 MAPK-dependent pathways in keratinocytes. Oncogene 19, 1596–1604. 10.1038/sj.onc.1203458.

Zhang, Z., Li, X., Yang, F., Chen, C., Liu, P., Ren, Y., Sun, P., Wang, Z., You, Y., Zeng, Y.X., and Li, X. (2021). DHHC9-mediated GLUT1 S-palmitoylation promotes glioblastoma glycolysis and tumorigenesis. Nat Commun 12, 5872. 10.1038/s41467-021-26180-4.

Zhang, Z., Zi, Z., Lee, E.E., Zhao, J., Contreras, D.C., South, A.P., Abel, E.D., Chong, B.F., Vandergriff, T., Hosler, G.A., et al. (2018). Differential glucose requirement in skin homeostasis and injury identifies a therapeutic target for psoriasis. Nat Med 24, 617–627. 10.1038/s41591-018-0003-0.

Zomer, H.D., and Trentin, A.G. (2018). Skin wound healing in humans and mice: Challenges in translational research. J Dermatol Sci 90, 3–12. 10.1016/j.jdermsci.2017.12.009.

